# Orphan nuclear receptors promote alternative lengthening of telomeres (ALT) through ALT-associated PML bodies

**DOI:** 10.1101/2022.03.30.485724

**Authors:** Venus Marie Gaela, Hsuan-Yu Hsia, Thomas Boudier, Liuh-Yow Chen

## Abstract

Alternative lengthening of telomeres (ALT) is a telomerase-independent telomere maintenance mechanism utilized by about 15% of cancers. Orphan nuclear receptors (NRs), such as COUP-TF1, COUP-TF2, EAR2, TR2, and TR4, associate with telomeres of ALT cells by binding to variant telomeric repeats. However, how these orphan NRs function in the ALT pathway remains to be characterized. Here, we have established an ALT-inducing cell model by tethering orphan NRs to telomeres in non-ALT BJ fibroblast cells. We demonstrate that recruitment of orphan NRs to telomeres is sufficient to initiate formation of ALT-associated promyelocytic leukemia nuclear bodies (APBs) and telomeric DNA synthesis at APBs. We found that the ability of orphan NRs to initiate APB formation and recombination is dependent on the orphan NR AF2 domain, the zinc-finger protein ZNF827, and PML protein. Depletion of orphan NRs in ALT cell lines reduced APB formation and telomeric DNA synthesis, confirming the role of orphan NRs in ALT cells. Furthermore, we found that ATRX/DAXX depletion, together with the telomeric localization of orphan NRs, induces APB formation, telomere clustering, and telomeric DNA synthesis more dramatically in non-ALT cells. Accordingly, we propose that these events in ALT, orphan NR recruitment to telomeres and ATRX/DAXX loss, operate in concert to activate the ALT pathway.

## INTRODUCTION

Telomeres are protective nucleoprotein structures at chromosome termini that prevent activation of DNA damage response and inhibit non-homologous end-joining. In humans, telomeres consist of 5’ TTAGGG 3’ DNA sequence repeats and telomere-binding proteins such as the shelterin complex (Moyzis et al., 1988; Palm & De Lange, 2008). Due to incomplete replication of the lagging DNA strand, telomeres shorten after each round of cell division, termed the “end-replication problem”. When telomeres become too short to maintain chromosome integrity and genome stability, cells enter apoptosis or senescence (Harley et al., 1990). Cancer cells overcome this problem by activating a telomere maintenance mechanism for their unlimited proliferation. In ∼85% of cancers, telomere length is maintained by reactivating telomerase. However, in ∼15% of cancers, particularly in tumors of mesenchymal or neuroepithelial origin, alternative lengthening of telomeres (ALT) acts as the mechanism for telomere maintenance. The ALT pathway is a telomerase-independent homologous recombination-based telomere maintenance mechanism that is thought to rely on break-induced replication (BIR) (Bryan et al., 1997; Dilley et al., 2016; Zhang et al., 2019). However, the initiating events that lead to activation of the ALT pathway remain unclear.

One hallmark of ALT cancers is the presence of ALT-associated promyelocytic leukemia nuclear bodies (APBs), which are defined as associations between promyelocytic leukemia nuclear bodies (PML-NBs) and telomeres (Yeager et al., 1999). PML-NBs are composed of the structural proteins PML and SP100, as well as associated proteins such as DAXX and SUMO. PML-NBs assemble via multivalent SUMO-SUMO-interacting motif (SIM) interactions, which may depend on liquid-liquid phase separation (LLPS) (Banani et al., 2016; Corpet et al., 2020). Previous studies have indicated that APB formation is promoted by replication stress, with DNA damage agents and loss of several replication stress response proteins increasing APBs in ALT cells (Cesare et al., 2009; Cox et al., 2016; Lu et al., 2019; Sobinoff et al., 2017). Sumoylation-mutant shelterin components TRF1 and TRF2 cannot form APBs, indicating that, like PML-NB formation, APB formation depends on sumoylation (Potts & Yu, 2007). Aside from their PML-NB components, APBs also contain recombination and repair proteins, supporting that APBs are sites for homologous recombination where a self-perpetuating loop of ALT activity may occur (Cho et al., 2014; Chung et al., 2012; Zhang et al., 2021). Knockdown or knockout of PML in an ALT cell line reduced recombination-mediated telomere synthesis, indicating that APBs are critical for telomere recombination (Loe et al., 2020; Zhang et al., 2019).

Genetic alterations may underlie ALT development. The most common mutations reported in ALT cancers and cell lines are mutations in genes encoding for α-thalassemia/mental retardation syndrome X-linked (ATRX) and death-domain-associated protein (DAXX) (Bower et al., 2012; Heaphy et al., 2011; Lovejoy et al., 2012; Schwartzentruber et al., 2012). Together, ATRX and DAXX form a histone chaperone complex that deposits histone variant H3.3 on telomeres and other heterochromatic regions (Drané et al., 2010; Elsässer et al., 2012; Goldberg et al., 2010; Lewis et al., 2010). ATRX and DAXX are known to be ALT suppressors, as evidenced by suppression of ALT upon ATRX re-expression (Clynes et al., 2015; Lovejoy et al., 2012; Napier et al., 2015). An *in vitro* cell immortalization study revealed that spontaneous loss of ATRX is an early event during ALT activation (Napier et al., 2015). Loss of ATRX stabilizes RNA:DNA hybrids and G-quadruplexes at telomeres, which induces replication stress that may promote ALT activation (Clynes et al., 2015; Flynn et al., 2015). Prolonged loss of ATRX or DAXX also leads to progressive telomere decompaction, which gradually induces telomere dysfunction (Li et al., 2019). Knockout of ATRX or DAXX also promotes several ALT-associated phenotypes, including APB formation (Li et al., 2019). However, loss of ATRX or DAXX is insufficient to activate ALT in non-ALT cell lines, implying that another genetic or epigenetic event cooperates with ATRX or DAXX loss to initiate the ALT pathway (Flynn et al., 2015; Lovejoy et al., 2012; Napier et al., 2015; O’Sullivan et al., 2014).

Telomeric localization of orphan nuclear receptors (NRs), including COUP-TF1, COUP-TF2, TR2, TR4, and EAR2, is a hallmark of ALT (Déjardin & Kingston, 2009). These orphan NRs belong to the nuclear hormone receptor superfamily and NR2C/F classes. COUP-TF2 is abundantly expressed during embryonic development and is involved in organogenesis, but its expression levels are reduced in adulthood (Pereira et al., 2000). Orphan NRs bind to telomeres of ALT cell lines and tumors via variant TCAGGG telomeric repeats, which are abundant in ALT telomeres but not in normal or telomerase-positive cells (Conomos et al., 2012). Several lines of evidence support a potential role for orphan NRs in the ALT pathway. Recruitment of the nucleosome remodeling and histone deacetylation (NuRD) complex to ALT telomeres by the zinc-finger protein ZNF827 is dependent on COUP-TF2 and TR4. Orphan NR-dependent recruitment of the NuRD-ZNF827 complex may cause remodeling of telomeric chromatin, rendering it more permissive to recombination (Conomos et al., 2014). COUP-TF2 and TR4 were also reported to interact directly with FANCD2, a protein involved in the Fanconi anemia DNA repair pathway, to induce a DNA damage response that contributes to the ALT pathway (Xu et al., 2019). However, the exact role of orphan NRs in ALT remains to be elucidated.

Here, we have recapitulated the ALT environment in non-ALT cells by artificially tethering orphan NRs to telomeres. Using this system, we elucidate the changes brought about by the recruitment of orphan NRs to telomeres, excluding the genetic and epigenetic differences in ALT cells. We demonstrate that tethering orphan NRs to telomeres is sufficient to induce ALT phenotypes, such as APB formation and telomere clustering. More importantly, recruitment of orphan NRs to telomeres initiates telomeric DNA synthesis at APBs in non-ALT cells. We have confirmed these roles of orphan NRs in ALT cell lines. Moreover, we added ATRX or DAXX depletion with recruitment of orphan NRs to telomeres and observed that these two events act cooperatively in ALT to promote APB formation, telomere clustering, and telomeric DNA synthesis even more dramatically. Taken together, we provide compelling evidence that orphan NR recruitment to telomeres triggers ALT via APB formation.

## RESULTS

### ERT2 tethering to telomeres induces APB formation and telomere clustering

To carefully characterize APB formation and telomere clustering, we utilized a ligand-dependent inducible system in BJ5TA^LT^ immortalized human fibroblasts cells. We utilized ERT2, the estrogen-receptor ligand-binding domain variant, belonging to the orphan NR superfamily and that shares a structural organization similar to orphan NRs. We tethered ERT2 to telomeres by fusing it with shelterin components TRF1 or TRF2 that bind directly to telomeres, generating ERT2-TRF1 and ERT2-TRF2, respectively. In the presence of the ERT2 ligand, 4-hydroxytamoxifen (4-OHT), both ERT2-TRF1 and ERT2-TRF2 accumulated in nuclei and localized at telomeres of BJ5TA^LT^ cells **(Fig. S1A)**. Using indirect immunofluorescence against PML coupled with telomere *in situ* hybridization, we observed that 4-OHT induced PML to localize at telomeres (indicative of APB formation) in BJ5TA^LT^ cells expressing ERT2-TRF1 and ERT2-TRF2, but not in cells expressing ERT2, TRF1, or TRF2 alone **(Fig. 1A-B)**. Apart from APB formation, we also detected a decrease in telomere number in BJ5TA^LT^ cells expressing ERT2-TRF1 and ERT2-TRF2 upon 4-OHT treatment, which could be indicative of telomere clustering **(Fig. 1C)**. Since our set-up is an inducible system, we investigated APB formation and telomere clustering in a time-dependent manner. Our time-course experiments show that APB formation and decrease in telomere number occurred 1 h after 4-OHT induction. The phenotypes became more pronounced over time and were sustained for at least 48 h after induction **(Fig. 1D-E)**.

**Figure 1.**
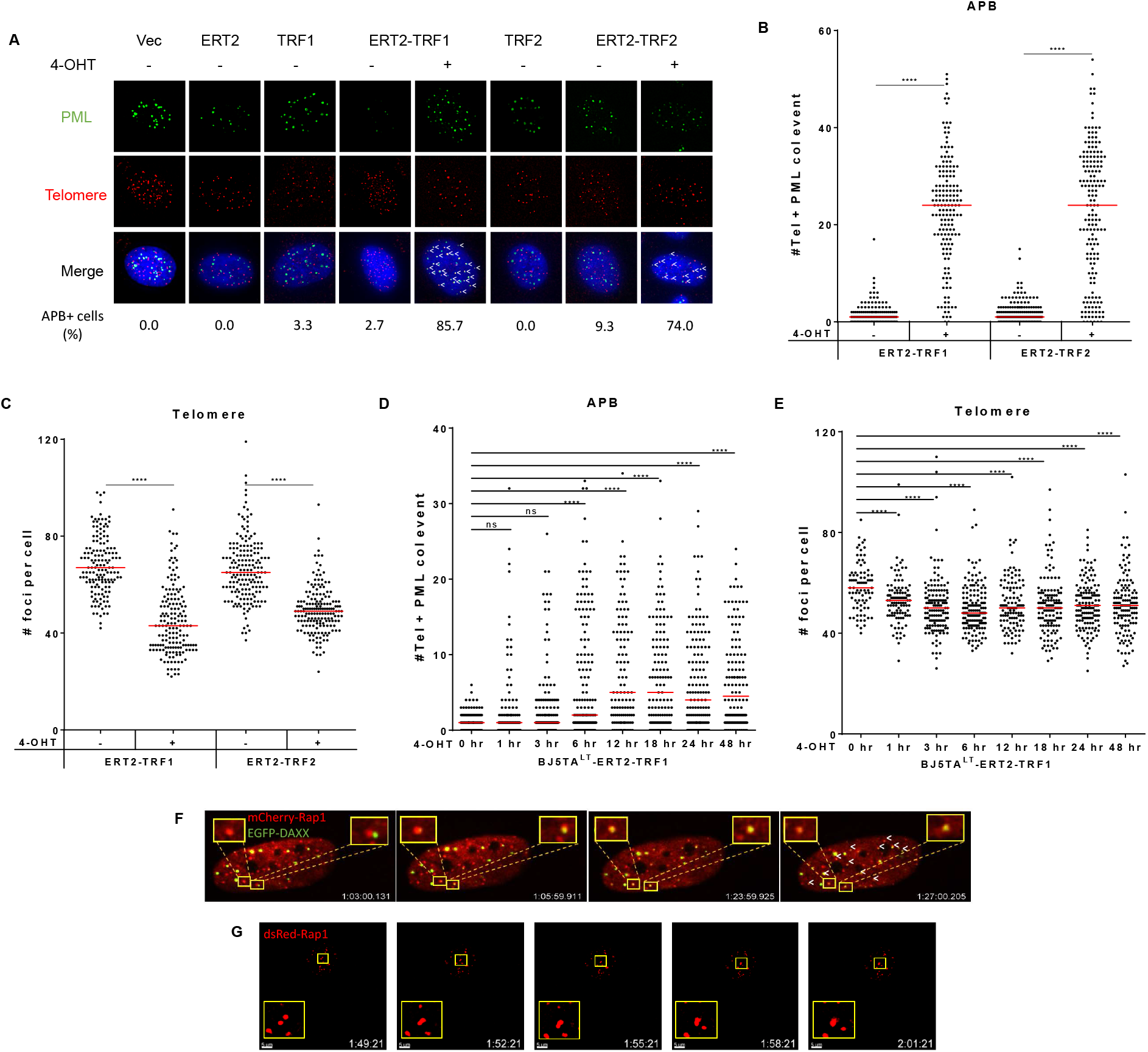
ERT2 tethering to telomeres induces APB formation and telomere clustering. **(A)** Representative images showing co-localization of PML and telomeres after 24 h of 4-OHT induction in BJ5TA^LT^-ERT2-TRF1 cells, but not in cells individually expressing ERT2, TRF1 or TRF2. PML was detected by immunofluorescence (IF), and telomeres were detected by FISH using the TelC PNA probe. Co-localization of PML (green) and telomeres (red) appears yellow. White arrows indicate APBs. APB+ cell percentage means more than five telomere+PML co-localization event. Quantification of APBs **(B)** and telomere numbers **(C)** in individual BJ5TA^LT^-ERT2-TRF1/ERT2-TRF2 cells after 24 h of 4-OHT induction. Quantification of APBs **(D)** and telomere numbers **(E)** in individual BJ5TA^LT^-ERT2-TRF1 cells (n >100) after 4-OHT time-course treatment (0, 1, 3, 6, 12, 18, 24, and 48 h). **(F)** Live-imaging of APBs. DAXX was fused with EGFP to visualize PML-NBs, and Rap1 was fused with mCherry to visualize telomeres. **(G)** Live-imaging of telomere clustering. DsRed-Rap1 indicates telomeres. Red lines represent median of duplicate experiments. ns p>0.05, *p<0.05 **p<0.01, ***p<0.001, ****p<0.0001, as determined by Mann-Whitney U test.

To understand the spatial dynamics of APB formation and telomere clustering, we established a live-imaging system using BJ5TA^LT^-ERT2-TRF2 cells. DAXX is another component of PML-NBs that co-localize with telomeres. We fused EGFP with DAXX to indicate PML-NBs and conjugated mCherry with Rap1 to indicate telomeres in BJ5TA^LT^-ERT2-TRF2 cells. We observed clear co-localization of EGFP-DAXX and mCherry-Rap1 foci 1-1.5 h after 4-OHT treatment **(Fig. 1F)**, indicating APB formation. We also visualized telomere dynamics in BJ5TA^LT^-ERT2-TRF2 using DsRed-Rap1 and observed clear telomeric clustering of two telomeres, giving rise to increased telomere volume and signal intensity 1-1.5 h after 4-OHT treatment **(Fig. 1G)**.

To further characterize telomere clustering, we developed a high-throughput quantification protocol for telomere signal intensity in Image J that allowed us to assess telomere clustering at different cell cycle stages. We imaged thousands of BJ5TA^LT^-ERT2-TRF1 cells before and after 4-OHT induction, and gated them to G1 and G2 phases of the cell cycle based on DNA content **(Fig. S2A)**. That strategy allowed us to compare telomere number and signal intensity in the same cell cycle stage. We observed a decrease in telomere number and an increase in telomere signal intensity in both the G1 and G2 phases upon 4-OHT induction **(Fig. S2B-C)**, indicating that the reduced number of telomeres is indeed an evidence of telomere clustering. By analyzing the telomeres in the same cell cycle stage, we ruled out the possibility that the observed decrease in telomere number was due to differences attributable to cell cycle stage. Furthermore, we observed a more dramatic decrease in telomere number and an increase in telomere signal intensity over time in the G2 phase after 4-OHT induction. It has been documented previously that APBs and telomere clustering are enhanced during the G2 phase and they decline during the transition to the G1 phase in ALT cells (Draskovic et al., 2009; Grobelny et al., 2000). Thus, our live-imaging experiment and quantification of telomere signal intensity provide clear evidence of APB formation and telomere clustering in BJ5TA^LT^-ERT2-TRF1/2 cells after induction, supporting that tethering ERT2 to telomeres via TRF1 or TRF2 triggers both phenotypes.

Aside from PML, SP100, and DAXX, PML-NBs also consist of sumoylation modifications with SUMO-interacting domains, which stabilize the spherical PML-SP100 shell (Chung et al., 2011). In analogy, it has been proposed that sumoylation of TRF1 and TRF2 by the MMS21-SMC5-SMC6 SUMO E3 ligase complex promotes recruitment of PML to telomeres (Potts & Yu, 2007). We examined whether active sumoylation is required for the APB formation induced by orphan NRs by using the SUMO E1 inhibitor, gingkolic acid. We observed a reduction in APB formation by inhibiting sumoylation in BJ5TA-ERT2-TRF2 cells **(Fig. S3A)**, but telomere clustering was unaffected **(Fig. S3B)**. Our data suggest that the APB formation induced by orphan NRs is also dependent on SUMO, consistent with the model proposed previously (Potts & Yu, 2007).

### Tethering of orphan NRs to telomeres induces APB formation and telomere clustering

Orphan NRs, including COUP-TF1, COUP-TF2, TR2, TR4, and EAR2, can bind to ALT telomeres via the variant TCAGGG or TGAGGG telomere repeats. However, the exact role of orphan NRs in ALT has been elusive. To investigate how orphan NRs contribute to ALT, we established an experimental model to recapitulate recruitment of orphan NRs in non-ALT BJ5TA^LT^ fibroblast cells, thereby accounting for genetic and epigenetic differences between ALT and non-ALT cells, such as telomere length heterogeneity, telomere damage, and mutations. We tethered the orphan NRs COUP-TF1, COUP-TF2, TR2, TR4, and EAR2 to BJ5TA^LT^ telomeres by fusing them to TRF1 and investigated for their ability to induce APB formation and telomere clustering. APB formation and telomere clustering occurred in BJ5TA^LT^ cells expressing TRF1-coupled COUP-TF1, COUP-TF2, TR2, TR4, or EAR2 **(Fig. 2A-C)**, demonstrating that recruitment of orphan NRs to telomeres induces both of these ALT phenotypes. We then determined which regions of orphan NRs are critical for their ability to induce APB formation and telomere clustering. Orphan NRs consist of a DNA-binding domain (DBD) and a ligand-binding domain (LBD). Since the LBD of orphan NRs has been reported previously to mediate interactions with co-activators or co-repressors (Kruse et al., 2008), we coupled the LBDs alone of COUP-TF1, COUP-TF2, TR2, TR4, and EAR2 with TRF1 in BJ5TA^LT^ cells and detected APB formation and telomere clustering in all BJ5TA^LT^ cells possessing orphan NR LBD-tethered telomeres **(Fig. 2D-F)**. Hence, recruitment of orphan NRs to telomeres is sufficient to induce APB formation and telomere clustering.

**Figure 2.**
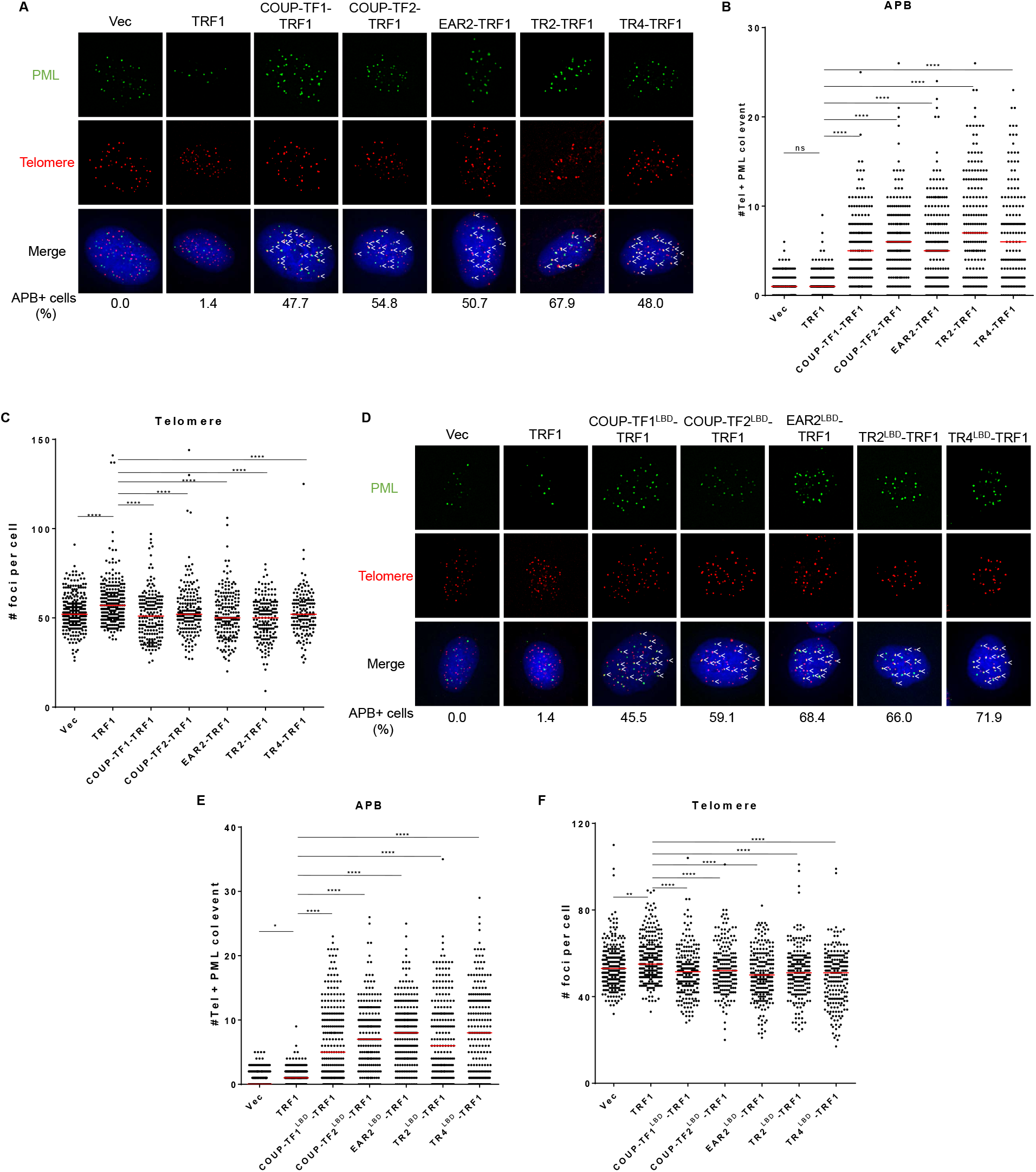
Tethering of orphan NRs to telomeres induces APB formation and telomere clustering. **(A)** Representative images showing PML and telomere co-localization in BJ5TA^LT^ cells expressing the full-length orphan NRs COUP-TF1/2, TR2/4, or EAR-2. PML was detected by IF, and telomeres were detected by FISH using the TelC PNA probe. Co-localization of PML (green) and telomeres (red) appears yellow. White arrows indicate APBs. APB+ cell percentage means more than five telomere+PML co-localization event. Quantification of APBs **(B)** and telomere numbers **(C)** in individual BJ5TA^LT^ cells (n>100) expressing the full-length orphan NRs COUP-TF1/2, TR2/4, or EAR-2. **(D)** Representative images showing co-localization of PML and telomeres in BJ5TA^LT^ cells expressing the LBD of orphan NRs COUP-TF1/2, TR2/4, or EAR-2. Quantification of APBs **(E)** and telomere numbers **(F)** in individual BJ5TA^LT^ cells (n>100) expressing LBDs of orphan NRs COUP-TF1/2, TR2/4, or EAR-2. Red lines represent median of duplicate experiments. ns p>0.05, *p<0.05 **p<0.01, ***p<0.001, ****p<0.0001, as determined by Mann-Whitney U test.

### The AF2 domain of orphan NRs is critical for APB formation and telomere clustering

Since tethering orphan NR LBDs to telomeres clearly triggered APB formation and telomere clustering, we wanted to identify which part of the LBD mediates the interaction. The LBDs of orphan NRs contain the activation function AF2 domain. It has been reported previously that the COUP-TF2 AF2 domain is responsible for the recruitment of co-activators (Kruse et al., 2008), which could conceivably induce APB formation and telomere clustering. We deleted the orphan NR AF2 domain of COUP-TF1 and COUP-TF2 and generated respective BJ5TA^LT^-COUP-TF1^LBDΔAF2^-TRF1 and COUP-TF2^LBD ΔAF2^-TRF1 cells, wherein APBs were reduced and telomere clustering was abolished upon AF2 domain deletion **(Fig. 3A-C)**. Our results indicate that the AF2 domain is critical for the ability of orphan NRs to promote APB formation and telomere clustering.

**Figure 3.**
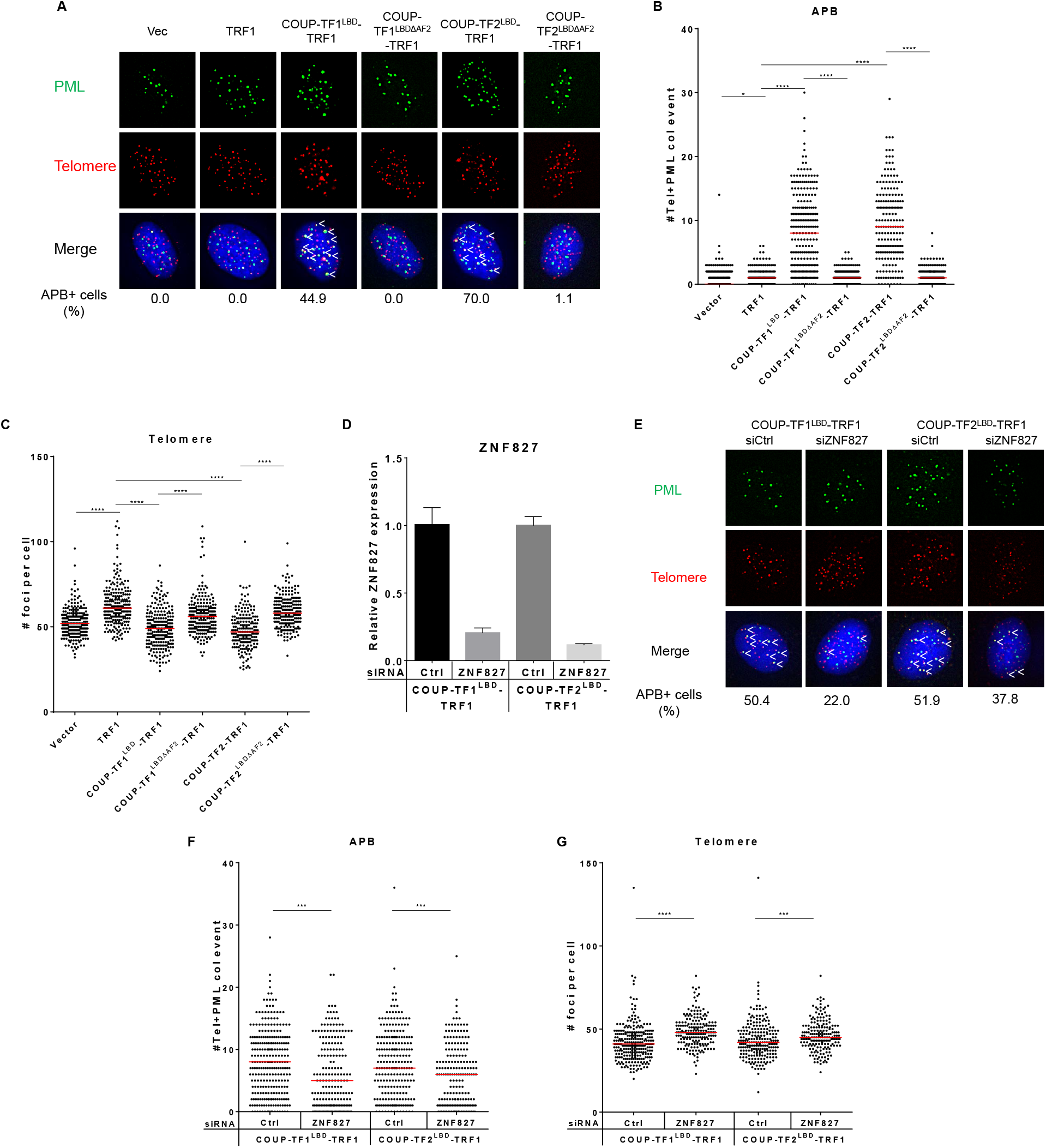
The AF2 domain of orphan NRs is critical for APB formation and telomere clustering. **(A)** Representative images showing loss of APBs in BJ5TA-COUPTF1/2^LBDΔAF2-TRF1^ cells. PML was detected by IF, and telomeres were detected by FISH using the TelC PNA probe. Co-localization of PML (green) and telomeres (red) appears yellow. White arrows indicate APBs. APB+ cell percentage means more than five telomere+PML co-localization event. Quantification of APBs **(B)** and telomere numbers **(C)** in individual BJ5TA^LT^ cells (n>100) expressing COUPTF1/2^LBD^-TRF1 or COUPTF1/2^LBDΔAF2-TRF1^. **(D)** Relative ZNF827 expression in BJ5TA^LT^ cells expressing COUPTF1/2^LBD^-TRF1, as measured by qPCR. **(E)** Representative images showing reduced APB formation in BJ5TA-COUP-TF1/2^LBD^TRF1 cells upon treatment with siRNAs against ZNF827. Quantification of APBs **(F)** and telomere numbers **(G)** in individual BJ5TA^LT^ cells (n>100) expressing COUPTF1/2^LBD^-TRF1 upon treatment with siRNAs against ZNF827. Red lines represent median of duplicate experiments. ns p>0.05, *p<0.05 **p<0.01, ***p<0.001, ****p<0.0001, as determined by Mann-Whitney U test.

Orphan NRs are proposed to be important for the recruitment of the ZNF827 zinc finger to ALT telomeres, which recruits the nucleosome remodeling and histone deacetylation (NuRD) complex to promote a chromatin environment suitable for homologous recombination (Conomos et al., 2014). We used siRNAs to deplete ZNF827 from BJ5TA-COUPTF1/2^LBD^-TRF1 cells, which resulted in fewer APBs and increased telomere number **(Fig. 3D-G)**, indicating that the ability of orphan NRs to induce APB formation and telomere clustering is dependent on ZNF827. Depletion of ZNF827 from WI38-VA13/2RA and IIICF/c cells was also reported to reduce APBs (Conomos et al., 2014), confirming that ZNF827 is indeed important for APB formation and telomere clustering.

Previous studies have indicated that sumoylation of TRF1 and TRF2 by the MMS21-SMC5-SMC6 SUMO E3 ligase complex promotes APB formation, followed by the recruitment of DNA repair and recombination agents to telomeres via SUMO-SIM interactions that trigger telomere compaction (Chung et al., 2011, 2012; Osterwald et al., 2015; Potts & Yu, 2007; Wu et al., 2003). We found that the recombination proteins NBS1, SMC5, and SMC6 also localize to telomeres in BJ5TA-COUPTF1/2^LBD^-TRF1 cells **(Fig. S4A)**. Intriguingly, we observed that all three of those proteins are already co-localized with PML-NBs in BJ5TA^LT^-Vector cells or before APB formation **(Fig. S4B)**, and their knockdown from BJ5TA^LT^-ERT2-TRF1 or BJ5TA^LT^-COUP-TF1/2^LBD^-TRF1 cells did not deplete either APB formation or telomere clustering **(Fig. S4C-H)**. Thus, none of these recombination proteins appear to be key players in recruiting PML to telomeres to initiate APB formation. However, we do not rule out the possibility that the SMC5/6 complex plays a role in the maintenance and stabilization of APBs. Nevertheless, our data support that tethering of orphan NRs to telomeres is sufficient to initiate APB formation and telomere clustering independently of the SMC5/6 complex. Accordingly, we propose that telomeric localization of orphan NRs directly initiates APB formation and telomere clustering via their AF2 domains and interactions with ZNF827, followed by the sumoylation of shelterin components by the SMC5/6 complex already present at PML-NBs and thereby further promotes APB formation and telomere clustering.

### Recruitment of orphan NRs to telomeres triggers telomeric DNA synthesis

We have shown that telomere tethering of orphan NRs induces ALT phenotypes, including APB formation and telomere clustering, in BJ fibroblast cells, but whether this initiates ALT telomere recombination remained unclear. To determine if recruitment of orphan NRs to the telomeres of BJ5TA^LT^ cells induced ALT activity, we utilized the ALT telomere DNA synthesis in APBs (ATSA) assay that enables direct detection of telomere replication in APBs during the G2 phase of the cell cycle (Zhang et al., 2019). Cells were synchronized in G2 phase with thymidine and CDK1 inhibitor treatments. Telomere extension was visualized by EdU (5-ethynyl-2’-deoxyuridine) incorporation, PML was detected by immunofluorescence, and telomeres were visualized by telomere fluorescence *in situ* hybridization. Some telomeres in BJ5TA^LT^ cells expressing COUP-TF1^LBD^-TRF1 and COUP-TF2^LBD^-TRF1 co-localized with EdU foci, indicative of DNA synthesis at telomeres **(Fig. 4A-B)**. Moreover, some of the EdU-associated telomeres also co-localized with PML foci, indicating telomere synthesis at APBs in the G2 phase **(Fig. 4A**,**C)**. Thus, we show, perhaps for the first time, that recruitment of orphan NRs to telomeres is sufficient to induce telomeric DNA synthesis at APBs in BJ5TA^LT^ non-ALT telomerase-positive cells.

**Figure 4.**
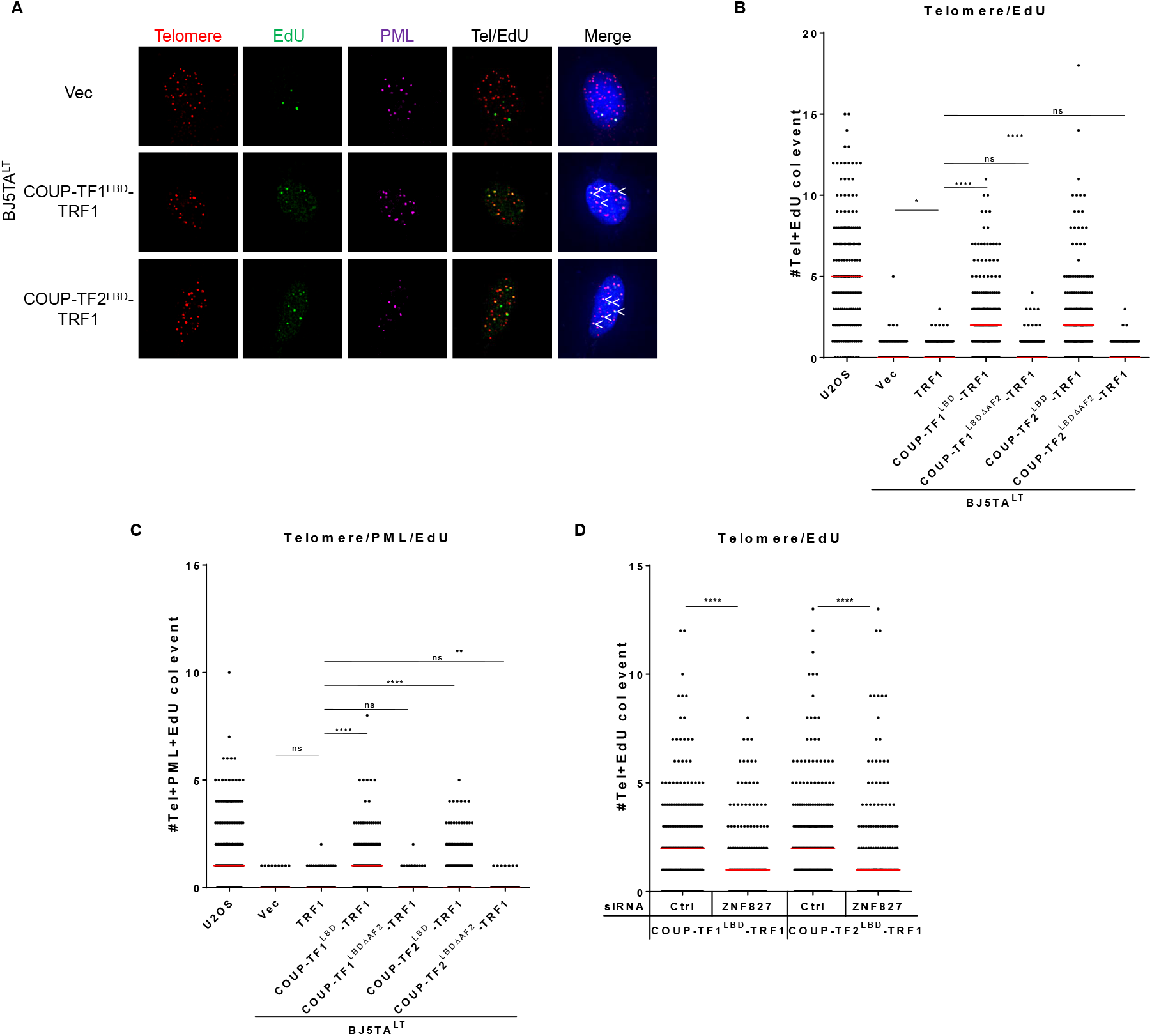
Recruitment of orphan NRs to telomeres triggers telomeric DNA synthesis. **(A)** Representative images showing EdU at telomeres and PML in BJ5TA^LT^-COUP-TF1/2^LBD^-TRF1 cells. EdU and PML were detected by IF, and telomeres were detected by FISH using the TelC PNA probe. Co-localization of EdU (green), PML (magenta), and telomeres (red) appears white. White arrows indicate telomeric DNA synthesis at APBs. **(B)** Quantification of telomere and EdU co-localization in BJ5TA^LT^-COUP-TF1/2^LBD^-TRF1 cells (n>100). **(C)** Quantification of telomere, PML, and EdU co-localization in BJ5TA^LT^-COUPTF1/2^LBD^-TRF1 cells (n>100). Red lines represent median of duplicate experiments. ns p>0.05, *p<0.05 **p<0.01, ***p<0.001, ****p<0.0001, as determined by Mann-Whitney U test. **(D)** Quantification of telomere and EdU co-localization in BJ5TA^LT^-COUP-TF1/2^LBD^-TRF1 cells (n>100) upon treatment with siRNAs against ZNF827. Cells were synchronized in G2 phase by thymidine and CDK1i treatments for 21 h and 12 h, respectively. Red lines represent median of duplicate experiments. ns p>0.05, *p<0.05 **p<0.01, ***p<0.001, ****p<0.0001, as determined by Mann-Whitney U test.

Since we had already uncovered that the orphan NR AF2 domain and ZNF827 are both critical for APB formation and telomere clustering, we wondered if they are also important for the initiation of telomeric DNA synthesis in BJ5TA^LT^ cells. Indeed, telomeric DNA synthesis at APBs was lost upon deleting the AF2 domain **(Fig. 4B-C)**. Similarly, upon depleting ZNF827, telomere DNA synthesis was reduced **(Fig. 4D)**. Our results demonstrate that orphan NR AF2 domain and ZNF827 are also critical to this process.

### PML is critical for orphan NR-induced telomeric DNA synthesis at APBs

Tethering of orphan NRs to telomeres induced both APB formation and telomere clustering in all of our experiments. To investigate the correlation between APB formation and telomere clustering, we generated PML knockout BJ5TA cells using the CRISPR/Cas9 technology. Guide RNAs targeting exons 1 to 3 of the *PML* gene generated an early stop codon leading to a knockout of all PML isoforms. Depletion of the PML protein was confirmed by immunofluorescence **(Fig. 5A)** and Western blot **(Fig. 5B)** analyses. We observed that telomere clustering still occurred in BJ5TA^LT^-ERT2-TRF1 and BJ5TA^LT^-COUP-TF1/2^LBD^-TRF1 cells upon PML knockout, suggesting that PML is not required for telomere clustering **(Fig. 5C-D)**. We also investigated whether PML is required for the ALT activity induced by orphan NRs and determined that telomeric DNA synthesis is suppressed in BJ5TA^LT^-COUPTF1/2^LBD^-TRF1 PML knockout cells **(Fig. 5E-F)**, indicating that PML is indeed critical for the ALT activity induced by the recruitment of orphan NRs to telomeres.

**Figure 5.**
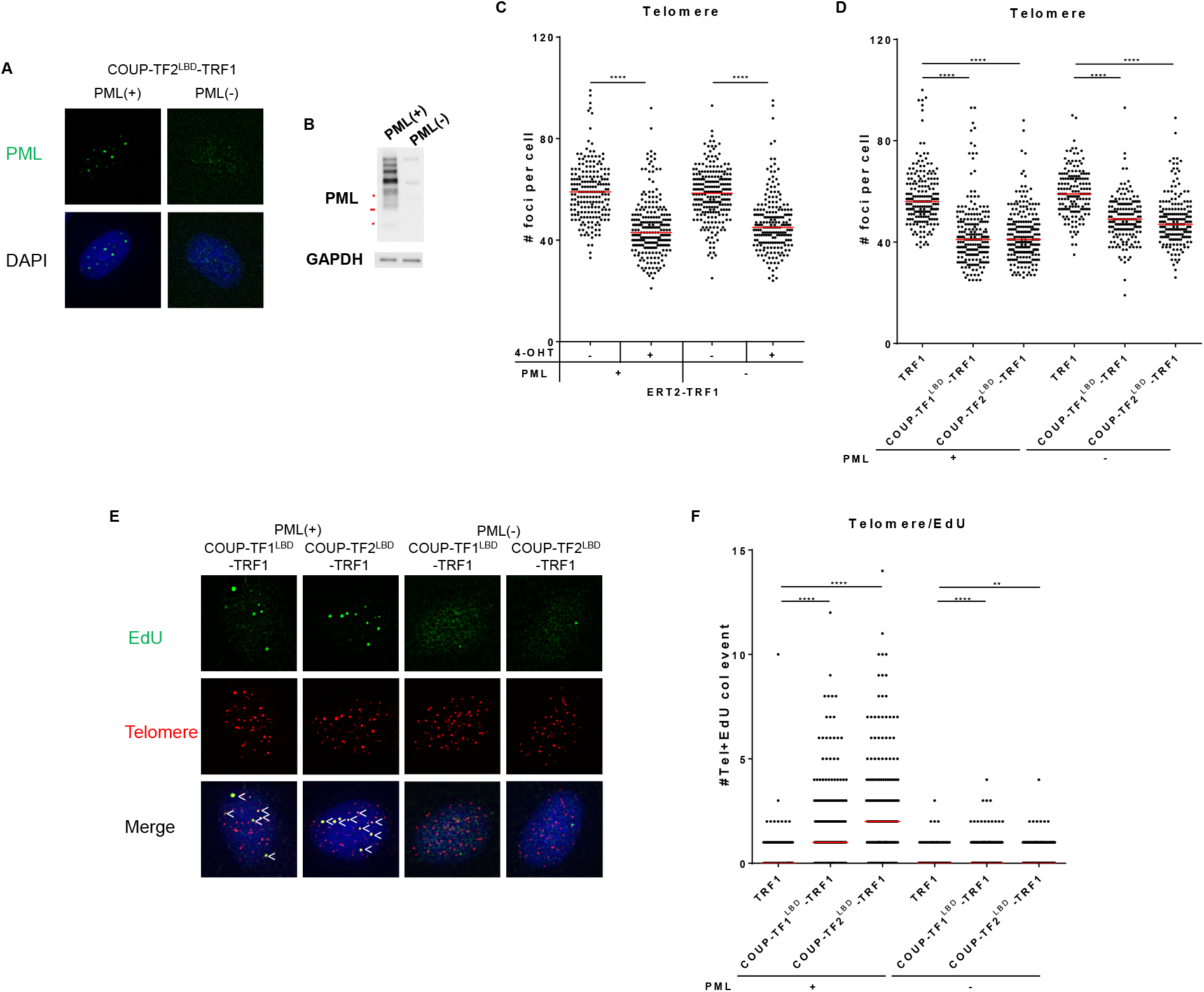
PML is critical for orphan NR-induced telomeric DNA synthesis at APBs. **(A)** Representative images of PML(+) and PML(-) BJ5TA-COUP-TF2^LBD^-TRF1 cells. PML was detected by IF. **(B)** Western blot showing PML protein expression in PML(+) and PML(-) BJ5TA cells. **(C-D)** Quantification of telomere number in PML(+) and PML(-) BJ5TA cells. **(E)** Representative images showing EdU at telomeres and PML in PML(+) and PML(-) BJ5TA^LT^ cells. EdU and PML were detected by IF, and telomeres were detected by FISH using the TelC PNA probe. Co-localization of EdU (green), PML (magenta), and telomeres (red) appears white. White arrows indicate telomeric DNA synthesis at APBs. **(F)** Quantification of telomere and EdU co-localization in PML(+) and PML(-) BJ5TA cells. Cells were synchronized in G2 phase by thymidine and CDK1i treatments for 21 h and 12 h, respectively. Red lines represent median of duplicate experiments. ns p>0.05, *p<0.05 **p<0.01, ***p<0.001, ****p<0.0001, as determined by Mann-Whitney U test.

### Orphan NRs mediate APB formation and telomere recombination in ALT cells

We have shown that recruitment of orphan NRs to telomeres in BJ5TA^LT^ non-ALT telomerase-positive cells is sufficient to induce APB formation, telomere clustering, and telomeric DNA synthesis at APBs. To determine if all these roles of orphan NRs are consistent in ALT cells, we depleted COUP-TF2 and TR4 from two ALT cell lines, i.e., U2OS and WI38-VA13/2RA. Single or combinatorial depletion of COUP-TF2 and TR4 by siRNAs decreased APB formation **(Fig. 6A-C)** and telomeric DNA synthesis **(Fig 6D-F)** in both those cell lines. Thus, the telomeric localization of orphan NRs is sufficient to promote telomeric DNA synthesis in both ALT and non-ALT cells.

**Figure 6.**
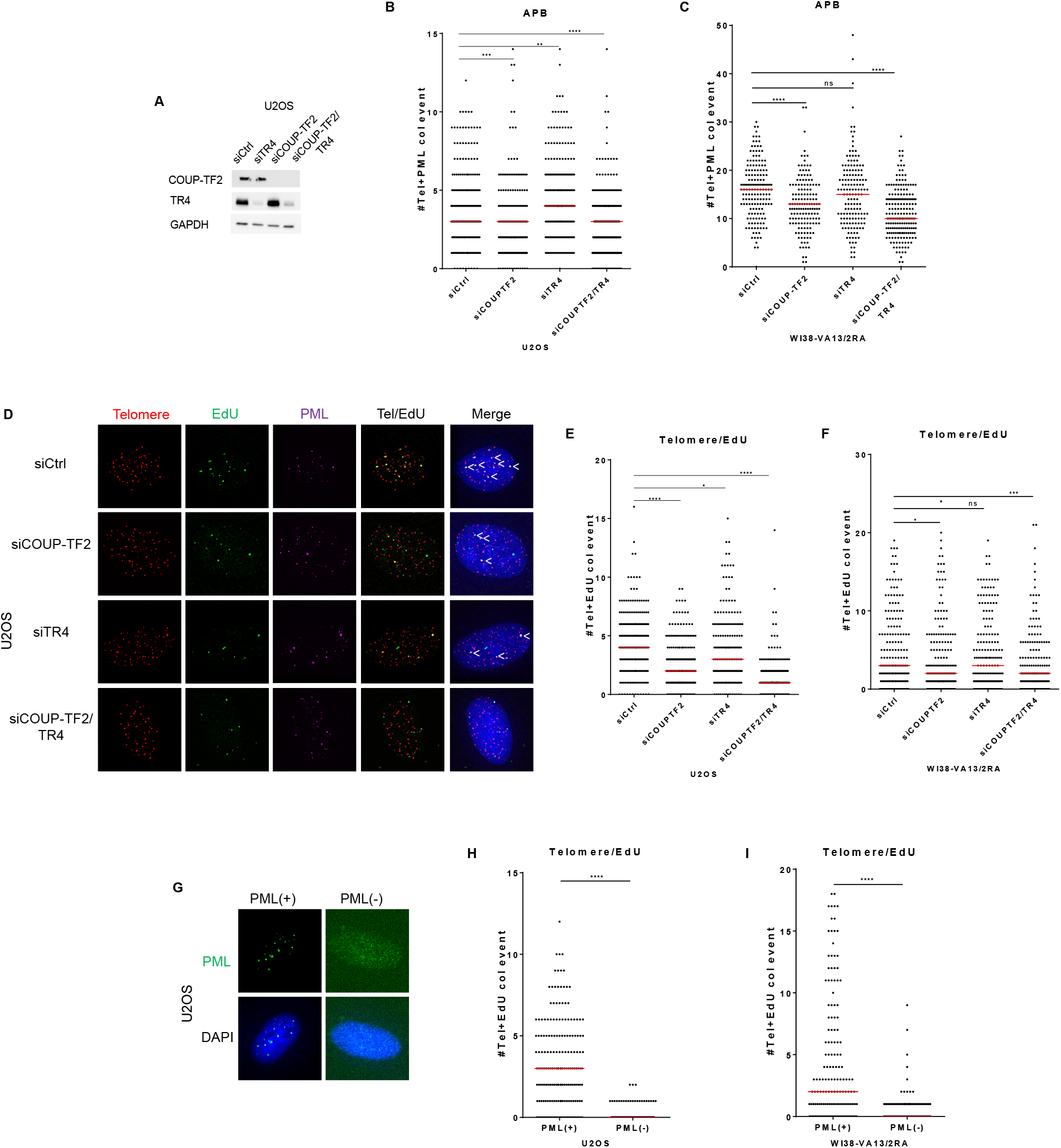
Orphan NRs mediate APB formation and telomere recombination in ALT cells. **(A)** Western blot showing COUP-TF2 and TR4 expression in U2OS cells. Quantification of APBs in U2OS **(B)** and WI38-VA13/2RA **(C)** cells upon treatment with siRNAs against COUP-TF2 or TR4. **(D)** Representative images showing reduced EdU signal at telomeres and PML levels in U2OS cells upon treatment with siRNAs against COUP-TF2 or TR4. **(E)** Quantification of telomere and EdU co-localization in U2OS cells upon treatment with siRNAs against COUP-TF2 or TR4. **(F)** Quantification of telomere, PML, and EdU co-localization in U2OS cells upon treatment with siRNAs against COUP-TF2 or TR4. **(G)** Representative images of PML(+) and PML(-) U2OS cells. **(H-I)** Quantification of telomere and EdU co-localization in PML(+) and PML(-) U2OS **(H)** and WI38-VA13/2RA **(I)** cells. Cells were synchronized in G2 phase bythymidine and CDK1i treatments for 21 h and 12 h, respectively. Red lines represent median of duplicate experiments. ns p>0.05, *p<0.05 **p<0.01, ***p<0.001, ****p<0.0001, as determined by Mann-Whitney U test.

To investigate whether PML is required for ALT activity in ALT cells, we also generated PML-knockout U2OS and WI38-VA13/2RA cells **(Fig. 6G)** and again observed that telomeric DNA synthesis was abolished in those cells **(Fig. 6H-I)**, similar to BJ5TA^LT^ cells and supporting findings by others (Loe et al., 2020; Zhang et al., 2019) that PML is critical for telomeric DNA synthesis.

### ATRX/DAXX depletion combined with telomeric recruitment of orphan NRs promotes ALT activity more dramatically

By tethering orphan NRs to telomeres in BJ5TA^LT^ fibroblast cells, we have shown that their recruitment to telomeres is sufficient to initiate ALT DNA synthesis in APBs. To recapitulate the ALT environment even more faithfully in BJ5TA^LT^ cells, we depleted them of the ATRX/DAXX/H3.3 deposition complex **(Fig. 7A)**. The ALT suppressors ATRX and DAXX form a histone chaperone complex that deposits histone variant H3.3 on telomeres (Elsässer et al., 2012; Goldberg et al., 2010; Heaphy et al., 2011; Lewis et al., 2010; Lovejoy et al., 2012). It was reported previously that depletion of ATRX/DAXX enhances ALT phenotypes, including APB formation (Li et al., 2019). Therefore, we hypothesized that a combination of these two ALT events, orphan NR recruitment to telomeres and ATRX/DAXX depletion, would result in much stronger ALT phenotypes in BJ5TA^LT^ cells. Indeed, we found that siRNA-mediated ATRX/DAXX depletion from BJ5TA^LT^-TRF1 control cells resulted in a slight increase in APB formation **(Fig. 7B)**, and ATRX/DAXX depletion from BJ5TA^LT^-COUP-TF1/2^LBD^-TRF1 cells induced a more prominent increase in APBs and telomere clustering **(Fig. 7B-C)**. We also determined that ATRX/DAXX depletion increased telomeric DNA synthesis at APBs in BJ5TA^LT^-TRF1 control cells **(Fig. 7D-F)** and enhanced telomeric DNA synthesis at APBs more markedly so in BJ5TA^LT^-COUP-TF1/2^LBD^-TRF1 cells **(Fig. 7D-F)**. Through this combinatorial mimicking of the ALT environment in BJ5TA^LT^ cells by recruiting orphan NRs to telomeres and depleting ATRX/DAXX, we have demonstrated that these two ALT events cooperate to initiate ALT telomeric DNA synthesis.

**Figure 7.**
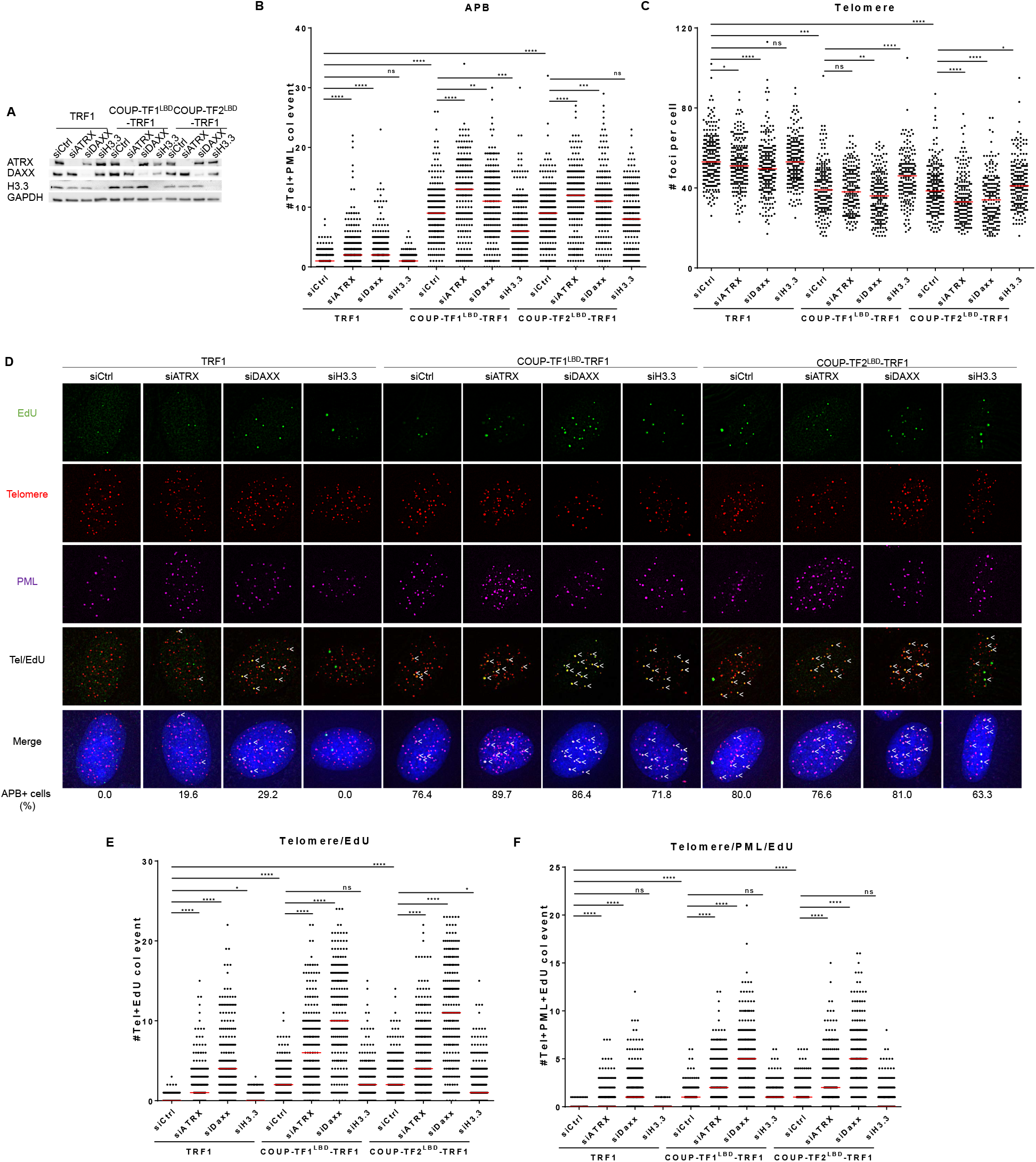
ATRX/DAXX depletion combined with orphan NR recruitment to telomeres promotes ALT activity more dramatically. **(A)** Western blot showing the loss of protein expression in BJ5TA^LT^-COUPTF1/2^LBD^-TRF1 cells upon treatment with siRNAs against ATRX, DAXX, or H3.3. Quantification of APBs **(B)** and telomere numbers **(C)** in BJ5TA^LT^-COUP-TF1/2^LBD^-TRF1 cells (n>100) upon treatment with siRNAs against ATRX, DAXX, or H3.3. **(D)** Representative images showing EdU at telomeres and PML in BJ5TA^LT^-COUP-TF1/2^LBD^-TRF1 cells upon treatment with siRNAs against ATRX, DAXX, or H3.3. EdU and PML were detected by IF, and telomeres were detected by FISH using the TelC PNA probe. Co-localization of EdU (green), PML (magenta), and telomeres (red) appears white. White arrows indicate telomeric DNA synthesis at APBs. APB+ cell percentage means more than five telomere+PML co-localization event. **(E)** Quantification of telomere and EdU co-localization in BJ5TA^LT^-COUP-TF1/2^LBD^-TRF1 cells (n>100) upon treatment with siRNAs against ATRX, DAXX, or H3.3. **(F)** Quantification of telomere, PML, and EdU co-localization in BJ5TA^LT^-COUP-TF1/2^LBD^-TRF1 cells upon treatment with siRNAs against ATRX, DAXX, or H3.3. Cells were synchronized in G2 phase bythymidine and CDK1i treatments for 21 h and 12 h, respectively. Red lines represent median of duplicate experiments. ns p>0.05, *p<0.05 **p<0.01, ***p<0.001, ****p<0.0001, as determined by Mann-Whitney U test.

## DISCUSSION

We have established a cell model system that induces ALT activity in non-ALT BJ5TA^LT^ human fibroblast cells by the recruitment of orphan NRs to telomeres. Tethering those orphan NRs to telomeres in non-ALT cells initiated ALT phenotypes such as APB formation and telomere clustering. More importantly, our results revealed for the first time that the telomeric localization of orphan NRs initiates telomeric DNA synthesis in non-ALT cells. These roles of orphan NRs are also consistent in ALT cell lines. Thus, our ALT-inducing model provides strong evidence that recruitment of orphan NRs to telomeres is an initial step in triggering ALT activity.

By artificially tethering orphan NRs to the telomeres of non-ALT cells, we could study the changes induced by telomeric recruitment of orphan NRs, representing a novel approach to determining how APBs are formed and how they facilitate ALT. Consistent with our results, previous studies have shown that depletion of the orphan NRs COUP-TF2 and TR4 from ALT cells abrogated some ALT phenotypes, such as formation of APBs and C-circles, as well as telomeric sister chromatid exchange (T-SCE) (Conomos et al., 2012; Déjardin & Kingston, 2009). However, it has also been postulated that orphan NRs only play an indirect role in ALT due to the absence of binding sites or variant repeats in some telomeres (Alhendi & Royle, 2020). In another study, variant repeat incorporation at telomeres using a mutant telomerase in a telomerase-positive cell line elicited several ALT hallmarks, but it did not activate inter-telomeric recombination-mediated copying (Conomos et al., 2014). Here, using the ATSA assay to visualize ALT telomere DNA synthesis in APBs, we have demonstrated that orphan NR recruitment to telomeres is sufficient to trigger telomeric DNA synthesis in non-ALT cells and to enhance telomeric DNA synthesis in ALT cells. Previous studies resorted to extreme situations to provide evidence of ALT recombination in non-ALT cells, such as via induction of high replication stress or a DNA damage response (Dilley et al., 2016; O’Sullivan et al., 2014). By circumventing those types of manipulation, our system provides clear evidence that recruitment of orphan NRs to telomeres is sufficient to initiate ALT DNA synthesis in APBs.

APBs are sites of homologous recombination and are critical for ALT activity (Yeager et al., 1999; Draskovic et al., 2009; Loe et al., 2019; Zhang et al., 2019). We and others (Loe et al., 2020; Zhang et al., 2019) have shown that depletion of PML from ALT cell lines abolishes telomeric DNA synthesis, consistent with our orphan NR-induced ALT cell model. It has been proposed that PML is important in ALT because it localizes the BLM-TOP3A-RMI (BTR) complex to ALT telomeres (Loe et al., 2020). In addition, all of our results support that APB induction is correlated with the initiation of telomeric DNA synthesis. We speculate that orphan NR recruitment to telomeres induces telomeric PML localization or APB formation, with this initial event further recruiting orphan NRs and more PML-NBs to telomeres to promote ALT activity.

Moreover, our findings indicate that PML is not necessary for telomere clustering. Although other studies have reported that PML depletion enhanced telomere signal intensity (Loe et al., 2020; Osterwald et al., 2015), at least one clone in the Loe et al. (2020) paper generated results similar to our PML-knockout clones, i.e., it still possessed PML-NB components at telomeres. Intriguingly, overexpression or knockdown of the orphan NRs COUP-TF2 and TR4 from ALT cells did not affect telomere clustering, nor did treating BJ5TA^LT^-ERT2-TRF1 cells with a sumoylation inhibitor. Hence, it is tempting to speculate that APB formation may be independent of telomere clustering. PML-NBs can associate with single or multiple telomeres, and APB formation may occur prior to telomere clustering. Accordingly, the nucleation event in initial APBs may drive recruitment of other recombination and repair factors to promote telomere clustering. Alternatively, telomeres may cluster in the absence of APBs. Whether or not telomere clustering directly induces telomeric DNA synthesis remains to be investigated.

It has been shown previously that ATRX depletion from mortal or immortal telomerase-positive cells is insufficient alone to activate ALT (Napier et al., 2015). However, we have demonstrated herein that combinatorial depletion of ATRX/DAXX and orphan NR recruitment to telomeres induced ALT DNA synthesis in non-ALT cells (Fig. 7). Based on that finding, we propose that orphan NR recruitment to telomeres and ATRX/DAXX loss represent two triggering events for ALT activation (Fig. 8). Recruitment of orphan NRs to telomeres may act in concert with the epigenetic effects of ATRX/DAXX loss to initiate ALT. ATRX/DAXX loss results in telomere decompaction, potentially eliciting dysfunctional telomere replication and activating homology-directed repair similar to ALT (Li et al., 2019). ATRX loss in non-ALT HeLa cells upregulates telomeric long noncoding RNAs called TERRA (telomeric repeat-containing RNA) in the G2/M phase of the cell cycle, mimicking the abundance of TERRA in the S and G2/M phases of ALT cells (Flynn et al., 2015). TERRA can generate RNA:DNA hybrids at telomeres (Flynn et al., 2015; Arora et al., 2014), which is a source of replication stress. Inhibiting TERRA transcription was shown previously to reduce DNA damage at telomeres, APB formation, and telomeric DNA synthesis in ALT cells (Silva et al., 2021). ATRX loss may also lead to stabilization of G-quadruplexes at telomeres (Clynes et al., 2015; Law et al., 2010), representing another source of replication stress that may activate ALT. ATRX depletion can also suppress resolution of the telomere cohesion that may enable telomere interaction leading to break-induced telomere recombination (Lovejoy et al., 2020; Ramamoorthy & Smith, 2015). The role of ATRX in promoting break-induced telomere recombination is reported to be DAXX-dependent (Lovejoy et al., 2020). Although DAXX depletion does not dramatically enhance APB formation (Fig. 7B, Li et al., 2019), it considerably promotes telomeric DNA synthesis (Fig. 7E-F). Despite not observing any significant changes in APB formation, telomere clustering, or telomeric DNA synthesis upon depleting histone H3.3, we cannot rule out the possibility that ATRX/DAXX loss results in decreased histone H3.3 deposition. Since histones display slow protein turnover, sufficient amounts of H3.3 may not have been depleted under our experimental conditions to induce that phenotype. Given that the histone deposition complex remains intact in H3.3-knockdown cells, the chromatin may not have been decompacted sufficiently to elicit phenotypes similar to those attributable to ATRX/DAXX loss. Establishing whether the telomere localization of orphan NRs induces transcription of TERRA and causes similar epigenetic effects to ATRX/DAXX loss warrants further investigation.

**Figure 8.**
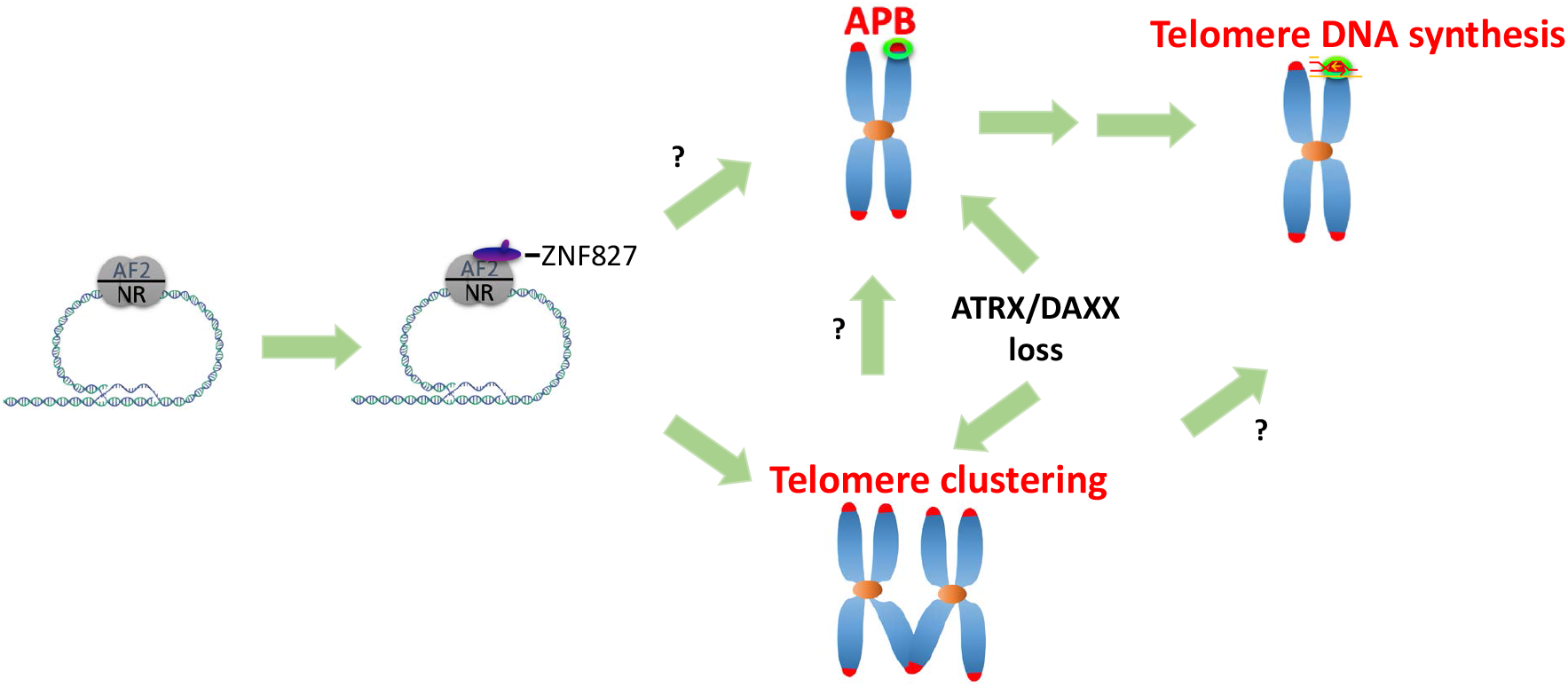
Model of orphan NR-induced ALT activation. Orphan NRs bind to ALT telomeres via variant repeats. The orphan NRs at telomeres recruit the NuRD-ZNF827 complex, which remodels the chromatin structure and allows APB formation and telomere clustering. The telomere localization of orphan NRs acts in concertwith ATRX/DAXX loss to promote APB formation and telomere clustering. Telomeric DNA synthesis can then be initiated at APBs.

How can orphan NRs initiate telomeric DNA synthesis? Orphan NRs bind to variant repeats of ALT telomeres and recruit the NuRD-ZNF827 complex, which has been proposed to affect chromatin structure by impeding shelterin binding and histone deacetylase-mediated hypoacetylation, leading to an environment suitable for APB formation and telomere recombination (Conomos et al., 2014). Recruitment of the NuRD-ZNF827 complex peaks at G2, i.e., the cell cycle phase when telomeric clustering is most prominent (Fig. S2) and when homologous recombination occurs. Indeed, we found that both the orphan NR AF2 domain and ZNF827 are critical for the induction of telomere recombination in non-ALT cells. There is some evidence that recruitment of ZNF827 to ALT telomeres may not be attributable to direct interaction, but instead may involve transcriptional suppression at telomeres (Yang et al., unpublished thesis). Recently, it was also reported that the LBDs of COUP-TF2 and TR4 directly interact with FANCD2 to induce a DNA damage response contributing to the ALT pathway (Xu et al., 2019). Our findings presented herein provide evidence supporting the model that orphan NRs associate with ALT telomeres and recruit NuRD-ZNF827 to create a chromatin environment that enables telomeric DNA synthesis. Overall, we have shown that recruitment of orphan NRs to telomeres initiates ALT activity in non-ALT cells via APB formation. Our results provide valuable information illuminating the mechanism underlying ALT cancer development that may be utilized to identify molecular targets for ALT cancer treatment.

## ACKNOWLEDGEMENTS

We thank L.-R. You of the Institute of Biochemistry and Molecular Biology at National Yang Ming University and S.-J. Chou of the Institute of Cellular and Organismic Biology at Academia Sinica for providing reagents, T.E. Chen for technical help, and S.-P. Chi of the Imaging Core in the Institute of Molecular Biology at Academia Sinica for help with data collection. Research in the laboratory of L.-Y. Chen was supported by grants from the Ministry of Science and Technology (110-2628-B-001-024) and Academia Sinica.

## AUTHOR CONTRIBUTIONS

V.M.G., H.-Y.H. and L.-Y.C. designed the study. V.M.G. performed most of the experiments, with assistance from H.-Y.H. T. B., V.M.G., and H.-Y.H. developed the protocol for image analysis. V.M.G and L.-Y.C. wrote the manuscript.

## MATERIALS AND METHODS

### Cell lines

Human fibroblast BJ5TA^LT^, human osteosarcoma U2OS, and human embryonic fibroblast 293T cells were cultured in DMEM (Gibco) supplemented with 10% fetal bovine serum (FBS) and 0.5% penicillin/streptomycin. SV40-transformed WI38/VA13-2RA fibroblasts were cultured in MEM (Gibco) supplemented with 10% FBS and 0.5% penicillin/streptomycin. All cells were grown at 37 °C with 5% CO2.

### Plasmids

ERT2-TRF1, ERT2-TRF2, COUP-TF1^LBD^-TRF1, COUP-TF2^LBD^-TRF1, EAR2^LBD^-TRF1, TR2^LBD^-TRF1, TR4^LBD^-TRF1, COUP-TF1^LBDΔAF2-TRF1^, and COUP-TF2^LBDΔAF2-TRF1^ were generated using the InFusion cloning method (Clontech). Vectors were digested with BamHI and incubated overnight at 37 °C (5 µg Vector, 10X CutSmart buffer 5 µl, BamHI 5 µl, H2O 35 µl). QIAquick PCR purification kit was used to purify the linearized vector. Target fragments were amplified by polymerase chain reaction (PCR), and PCR products were treated with DpnI before purification. The 5X InFusion HD Enzyme Premix, linearized vector, purified PCR fragment, and dH2O were mixed and incubated at 50 °C for 15 min. Transformation was conducted using stbl3 (ECOS™-competent cells) with ampicillin selection and overnight incubation at 37 °C. Single clones were selected and amplified by liquid culture. Plasmids were collected using QIAprep Spin Miniprep Kit and were checked by sequencing and restriction digestion. Retroviral transduction was performed to stably express the plasmids in cells.

### RNA interference

siRNA transfections were done by reverse transfection with Lipofectamine RNAiMax (Invitrogen). All siRNAs were transfected at 25 uM as per the manufacturer’s recommendations. For synchronization using thymidine and CDK1 inhibitor (CDK1i, RO-3306), cells were treated with thymidine 48h post transfection.

### Generation of CRISPR PML-knockout cell lines

BJ5TA^LT^ cells stably expressing doxycycline-inducible Cas9 were first generated by lentiviral transduction. The stable cells were transfected with PML guide RNAs using Lipofectamine RNAiMAX, according to the manufacturer’s instructions (Invitrogen, USA), simultaneous with doxycycline treatment (50 ng/ml) for 3 days. For U2OS and WI38-VA13/2RA PML-knockout cells, U2OS and WI38-VA13/2RA doxycycline-inducible Cas9 cells were transduced with PML guide RNAs, and then treated with doxycycline (50 ng/ml) for 3 days. Single clones were isolated using limiting dilution. Knockout efficiency was checked by immunofluorescence and Western blot. Retroviral transduction was then used to stably express TRF1, ERT2-TRF1, COUP-TF1^LBD^-TRF1, or COUP-TF2^LBD^-TRF1 in PML(+) and PML(-) BJ5TA cells.

### Detection of telomeric DNA synthesis

To synchronize cells at G2 phase, cells were treated with 2 mM thymidine for 21 h, released into fresh medium for 4 h, and then treated with 15 mM CDK1i for 12 h. To visualize DNA synthesis, cells were incubated with 20 mM EdU for 3 h. EdU was labeled with the fluorescent dye picolyl azide via click-it reaction (Invitrogen).

### Immunofluorescence (IF) and fluorescence *in situ* hybridization (FISH)

Cells were seeded onto coverslips inserted into 12-well plates. Cells were fixed with 4% paraformaldehyde for 10 min and permeabilized with 0.5% Triton-X-100 (diluted in phosphate buffered saline, PBS) for 5 min. Cells were blocked with 2% FBS (diluted in PBS) for 30 min. Primary antibodies (1:200) and secondary antibodies (1:800) were diluted in 2% FBS, and cells were incubated for 1 h each. Cells were fixed again with 4% paraformaldehyde for 10 min. Serial dehydration in 70% EtOH, 95% EtOH, and 100% EtOH was conducted for 5 min at each concentration, before air-drying the coverslips. Hybridization mix (10 mM Tris-HCl pH 7.5, 70% formamide, 10% blocking reagent, 0.25 µM telomere probe) was then placed on the slide. Denaturation at 80 °C on a heat-plate for 3 min was done, followed by overnight hybridization at room temperature. Cells were washed twice with wash buffer A (70% formamide, 10 mM Tris-HCl pH7.5), and then three times with wash buffer B (10 mM Tris-HCL pH 7.5, 4 M NaCl, 20% Tween-20). DAPI (Life Technology) was used to stain nuclei. Serial alcohol dehydration was performed before again air-drying the coverslips and then mounting on a slide with Prolong Gold Antifade Mountant (Life Technology).

### Image acquisition and quantification

Fluorescence 3D images were acquired using a GE Healthcare DeltaVision Deconvolution microscope. Images were taken with 0.2 µm spacing for a total of 5 µm. The images were analyzed using softWoRx 5.5.1 for deconvolution and ImageJ/FIJI for quantification. First, a maximum intensity projection was applied to visualize the acquired Z-stack as a 2D-image. Then, the “tophat” filter was used for background subtraction. The “find maxima” command was used to identify the local maxima of signal intensity in each image, wherein the results are the segmented objects identified for each telomere or each PML-NB. Finally, overlap of objects in telomere and PML-NB images were recognized as APBs. To automatically quantify telomere and APB numbers, we utilized the ImageJ Macro language (IJM), which is a scripting language built into ImageJ.

### Live-imaging

Cells were seeded on 22 × 22-mm glass coverslips in single wells of a 6-well plate. For imaging, coverslips were mounted in magnetic chambers with cells maintained in DMEM supplemented with 10% FBS and 1% penicillin/streptomycin at 37 °C on a heated stage in an environmental chamber. Images were acquired using a GE Healthcare DeltaVision Deconvolution microscope. 4-OHT was added after capturing the first frame. For APB formation, images were taken at 3-min time intervals for 1.5 h for both the GFP and mCherry channels. For telomere clustering, images were taken at 30-sec intervals from frames 2-121, and at 3-min intervals thereafter for a total of 7 h for the DsRed channel.

### Statistical analyses

GraphPad Prism 6, Matlab, and Microsoft Excel were used to generate tables and graphs. All statistical analyses were performed using two-tailed Mann-Whitney tests.

## SUPPLEMENTARY FIGURE LEGENDS

**Figure S1.**
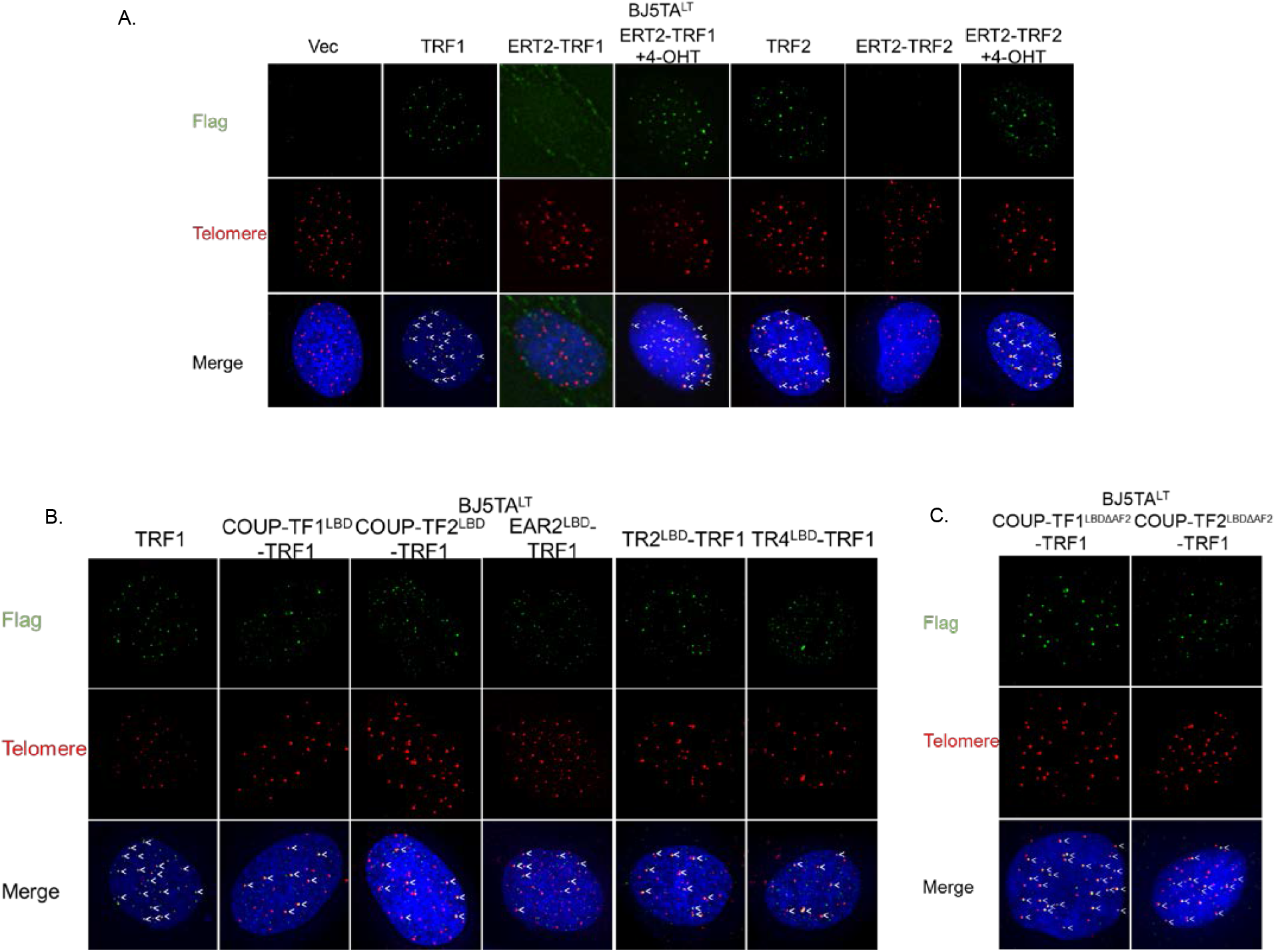
Orphan NRs tether to telomeres. **(A)** Representative images showing co-localization of Flag-tagged ERT2-TRF1 and ERT2-TRF2 fusion proteins with telomeres in BJ5TA^LT^ cells upon addition of 4-OHT. **(B)** Representative images showing co-localization of Flag-tagged COUP-TF1^LBD^-TRF1, COUP-TF2^LBD^-TRF1, EAR2^LBD^-TRF1, TR2^LBD^-TRF1, and TR4^LBD^-TRF1 fusion proteins with telomeres in BJ5TA^LT^ cells. **(C)** Representative images showing co-localization of Flag-tagged COUP-TF1^LBDΔAF2-TRF1^ and COUP-TF2^LBDΔAF2-TRF1^ fusion proteins with telomeres. The Flag tag was detected by immunofluorescence (IF), and telomeres were detected by FISH using the TelC PNA probe. Co-localization of Flag (green) and telomeres (red) appears yellow. White arrows indicate co-localization of Flag-tagged proteins and telomeres.

**Figure S2.**
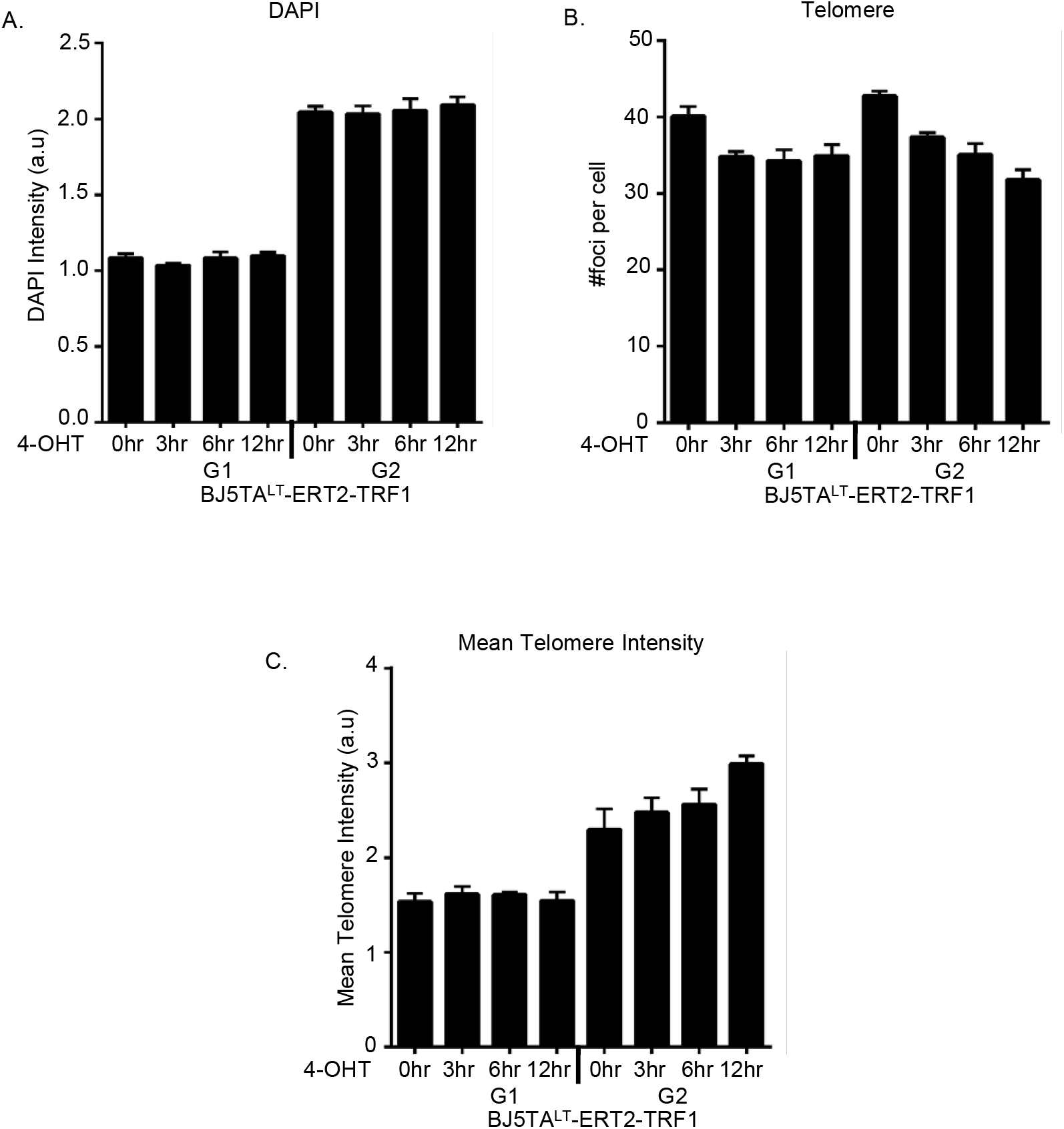
Telomere clustering is more pronounced in the G2 phase of the cell cycle. **(A-C)** Quantification of DNA content, telomere number, and telomere intensity in BJ5TA^LT^-ERT2-TRF1 cells at different cell cycle stages. Cells in the G1 and G2 phases were gated according to DAPI signal. The average of three replicates is shown (n>500 for each replicate), with the error bar representing SD.

**Figure S3.**
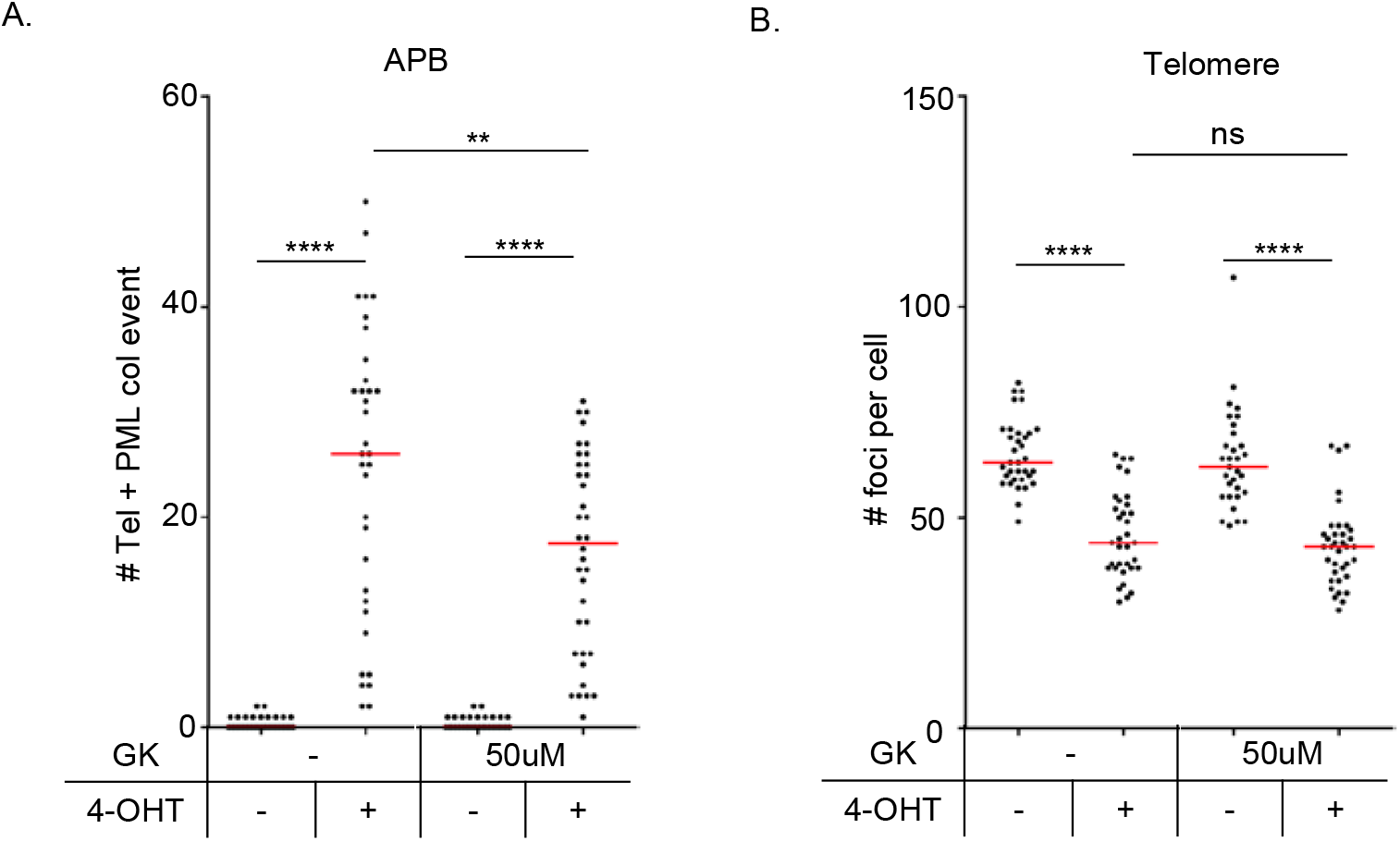
APB formation is reduced upon sumoylation inhibition. Quantification of APBs **(A)** and telomere numbers **(B)** in BJ5TA^LT^-ERT2-TRF2 cells after 24 h of treatment with 50 or 100 µM gingkolic acid plus 24 h 4-OHT induction. Red lines represent median. ns p>0.05, *p<0.05 **p<0.01, ***p<0.001, ****p<0.0001, as determined by Mann-Whitney U test.

**Figure S4.**
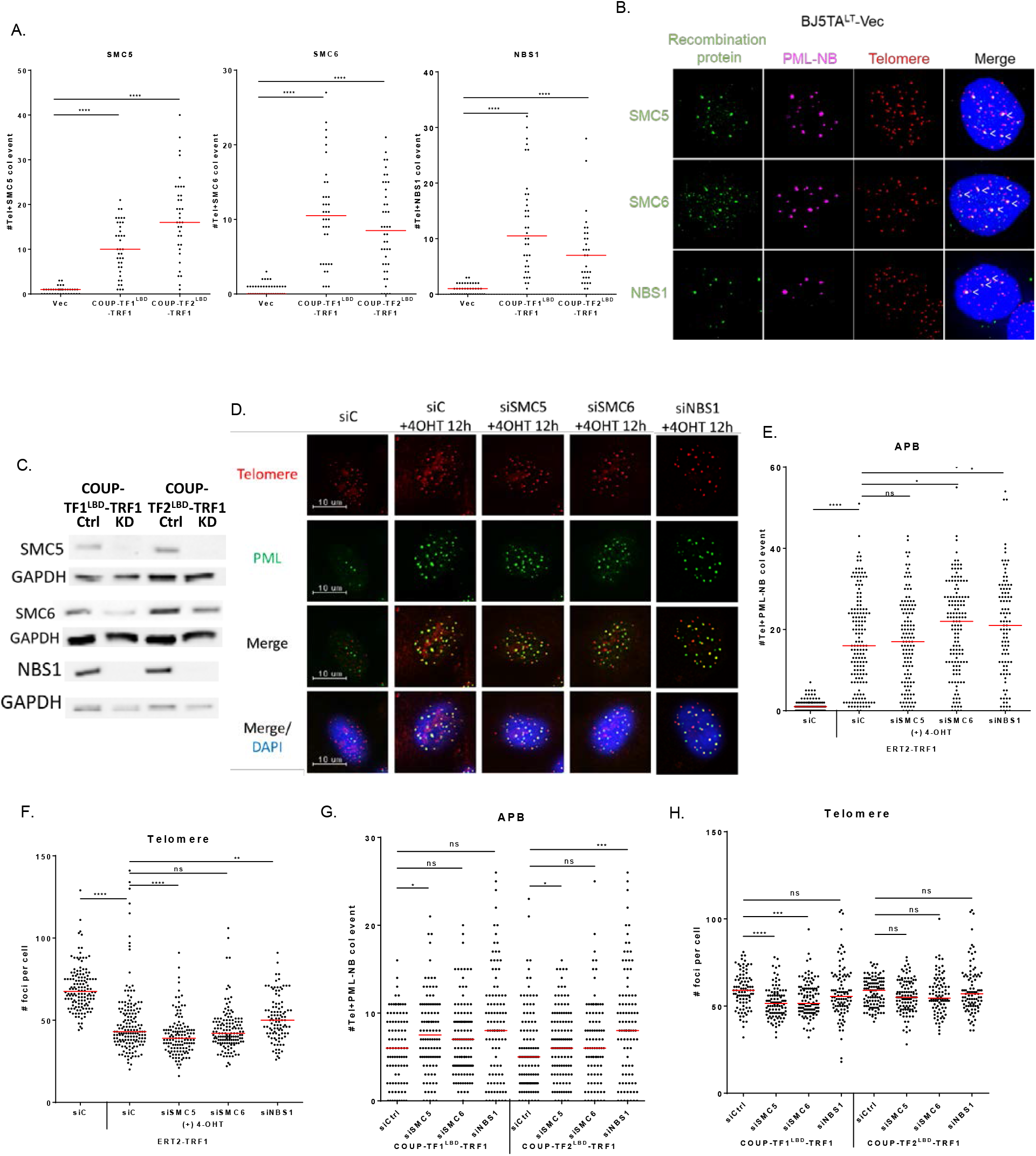
Recombination proteins localize to PML-NBs before APB formation. **(A)** Quantification of the co-localization of recombination proteins NBS1, SMC5, and SMC6 with telomeres in BJ5TA^LT^-COUPTF1/2^LBD^-TRF1 cells. **(B)** Representative images showing co-localization of PML-NBs and recombination proteins before APB formation in BJ5TA^LT^-Vec cells. Recombination proteins (SMC5/6 and NBS1) and PML/DAXX were detected by IF, and telomeres were detected by FISH using the TelC PNA probe. Co-localization of recombination proteins (green) and PML/DAXX (magenta) appears white. White arrows indicate co-localization of recombination proteins and PML-NBs in BJ5TA cells. **(C)** Western blot showing reduced protein expression in BJ5TA^LT^-COUP-TF1/2^LBD^-TRF1 cells upon treatment with siRNAs against SMC5/6 or NBS1. **(D)** Representative images showing co-localization of PML and telomeres in BJ5TA^LT^-ERT2-TRF1 cells even after treatment with siRNAs against SMC5/6 or NBS1. Quantification of APBs **(E)** and telomere numbers **(F)** in individual BJ5TA^LT^-ERT2-TRF1 cells (n>100) upon treatment with siRNAs against SMC5/6 or NBS1. Quantification of APBs **(G)** and telomere numbers **(H)** in individual BJ5TA^LT^-COUP-TF1/2^LBD^-TRF1 cells (n>100) upon treatment with siRNAs against SMC5/6 or NBS1. Red lines indicate median. ns p>0.05, *p<0.05 **p<0.01, ***p<0.001, ****p<0.0001, as determined by Mann-Whitney U test.

